# Stimulus-specificity of surround-induced responses in primary visual cortex

**DOI:** 10.1101/2024.06.03.597080

**Authors:** Nisa Cuevas, Boris Sotomayor-Gómez, Athanasia Tzanou, Irene Onorato, Brian Rummell, Cem Uran, Ana Broggini, Martin Vinck

## Abstract

Recent work suggests that stimuli in the surround can drive V1 neurons even without direct visual input to the classical receptive field (RF). These surround-induced responses may represent a prediction of the occluded stimulus, a prediction error, or alternatively, a representation of the gray patch covering the RF. Using Neuropixels recordings in mouse V1, we found that a distal surround stimulus increased V1 firing rates for gray patches up to 90° in diameter, while LGN firing rates decreased for the same stimuli. These responses occurred across a wide range of conditions: they were elicited by both moving and stationary surround stimuli, did not require spatial continuity or motion coherence, and persisted even for large gray patches (90°) where there was no mismatch between the classical RF stimulus (∼20°) and the near surround. They also emerged when the gray patch appeared as a salient object against a uniform black or white background. Additionally, response magnitudes and latencies were highly similar for black/white uniform surface stimuli on a gray background, with latencies increasing with the gray-patch diameter. These findings are difficult to reconcile with the predictive coding interpretation and fit best with the hypothesis that surround-induced responses reflect the representation of the uniform surface itself and may thereby contribute to image segmentation processes.

## Introduction

A characteristic feature of cortical circuits is the integration of feedforward afferent inputs with horizontal and top-down feedback (***Angelucci et al., 2002, 2017; Gilbert and Wiesel, 1990; Vezoli et al., 2021***). This integration may account for the observation that sensory responses in the neocortex exhibit a systematic dependence on the spatiotemporal context and cognitive factors like attention, working memory, etc. (***Gazzaley and Nobre, 2012; Quak et al., 2015; Pasternak and Greenlee, 2005; De Lange et al., 2018***). While feedforward influences are commonly conceptualized as “driving”, feedback has been characterized as “modulatory”, i.e. strengthening or diminishing the feedforward responses, without driving neural responses in the absence of feedforward inputs (***Vezoli et al., 2021; Cavanaugh et al., 2002b; Self et al., 2013***). Receptive fields in primary visual cortex (V1) are usually conceptualized along these lines, with a classic receptive field (RF) that accounts for evoked sensory responses and an extra-classical surround that allows for further modulation of sensory responses by spatial context (***Maffei and Fiorentini, 1976; Allman et al., 1985; Angelucci et al., 2002, 2017***). However, recent studies in mice challenge the idea that the surround acts in a purely modulatory manner (although we note that there are some inconsistencies between primate studies ***Slllito et al. 1995; Cavanaugh et al. 2002a; Gieselmann and Thiele 2008***, see Discussion).

Specifically, ***Keller et al. (2020***) report that in mouse V1, firing rates are higher when a drifting grating is partially occluded by a gray mask over the classical RF compared to an unoccluded fullfield drifting grating. ***Keller et al. (2020***) describe this phenomenon as an inverse (or second) RF, emphasizing the driving influence of the surround. Optogenetic manipulation further suggests that the reported increase in V1 firing rates for the gray-patch condition depends on top-down feedback from secondary visual cortical areas (***Keller et al., 2020***). These findings from ***Keller et al. (2020***) raise two major questions:

First, we argue that it is still uncertain whether stimuli presented in the surround alone can independently drive neural responses, which we refer to as a “surround-induced response”. In particular, the strongest increase in firing rates in (***Keller et al., 2020***) occurred for gray patches around 15°diameter. For such a small gray patch, part of the grating stimulus may still activate the classical RF, particularly given that neurons with RFs up to 10° from the patch center were included (***Keller et al., 2020***, see Discussion). In other words, the gray patch may introduce a new stimulus (grating/gray) within the RF, increasing firing rates because of a mismatch with the surround. The results for large surrounds, where any remaining stimulation of the classic RF can be excluded, were not systematically investigated and appear to be inconclusive (***Keller et al., 2020***). To address this issue, it is necessary to systematically investigate V1 responses for large-diameter gray patches where bottom-up sensory stimulation can be excluded.

Second, the stimulus-dependence of surround-induced responses remains to be systematically investigated, which is necessary to determine the computational mechanisms and functional significance of surround-induced responses. In particular, we wish to contrast two interpretations of surround-induced responses: (i) One interpretation is that surround-induced responses result from predictive processing (***Rao and Ballard, 1999***) (“predictive processing hypothesis”). In this interpretation, the inverse RF may reflect either an omission signal, resulting from the absence of a predicted input, or a prediction signal of the occluded content (***Muckli et al., 2015; Keller et al., 2020***). These predictive processing explanations entail that the properties of the surround stimulus should be a critical factor. One would expect that prediction error or predictive fill-in signals will be boosted when the visual system can infer that there is a stimulus behind the gray patch, which thereby acts as an “occluder”. We reckoned that the inference of a stimulus behind the gray mask would be facilitated when the surround stimulus appears to be moving behind the occluder. Furthermore, we reasoned that prediction signals should depend on the spatial or motion coherence of the surround stimulus, as a coherent surround leads to interpolation and increases the precision of predictions. In this study, we tested these predictions by comparing moving *vs*. stationary stimuli and manipulating the spatial or movement coherence of the surround stimulus.

(ii) An alternative explanation for surround-induced responses is that these responses reflect the representation of the gray patch itself and relate to segmentation processes (“segmentation hypothesis”). In macaque V1, ***Zweig et al. (2015***) have shown that responses to black or white uniform surfaces have longer latencies at the stimulus center compared to the edge and show a systematic increase in response latency with the size of the surface stimulus. ***Zweig et al. (2015***) suggests that this increase in latency reflects the perceptual inference of the uniform surface information, requiring information transfer from the surface edge towards the center of the surface. The rationale here is that at the center of an achromatic surface stimulus, there is no intrinsic signal of the surface properties and that these properties are inferred by using information from the surface’s edge ***Zweig et al. (2015***). In this scenario, presenting a stimulus in the surround may create a transient activation around the edge that leads to a transient enhancement of the representation of the gray surface itself (***Peter et al., 2019***). This “segmentation” interpretation would entail that the surround-induced responses for gray masks should be very similar to the case of black/white center stimuli with a uniform gray surround. Another prediction is that there should be a systematic increase in the response latency with the size of the gray patch (***Zweig et al., 2015***). Finally, the “segmentation hypothesis” does not require spatial or motion coherence of the surround stimulus, because a gray patch will be visible both for a spatially coherent and incoherent surround.

In the present study, we recorded V1 and lateral geniculate nucleus (LGN) neurons using Neuropixels in awake mice. We demonstrate that neural responses in V1 can increase with stimulation of the distal surround, up to 90°diameter, while LGN firing rates decrease for the same stimuli. Based on these observations, we performed a detailed investigation of the neural responses to distal surround stimuli with a large gray patch covering the RF. We systematically investigated the dependence of the neural response on the properties of the surrounding stimuli using single- unit and population decoding analyses. We presented four kinds types of stimuli: (1) stationary and drifting gratings that were spatially continuous; (2) surround stimuli that were divided into two drifting gratings that lacked motion coherence or spatially discontinuous static gratings; (3) noisy textures; and (4) black/white surface stimuli on a gray surround, or gray surface stimuli in a black/white surround.

## Results

### Responses to occluded grating stimuli

We used Neuropixel probes to record neuronal activity across all layers of V1 in head-fixed mice placed on a running disk (Figure 1a,b; see Methods). Visual stimuli were centered on the RFs of the recorded neurons (Figure 1c, see Methods). We included only single units into the analysis that met several criteria in terms of visual responsiveness and a significant RF with a center within 10° (absolute) distance to the stimulus center of the grating.

**Figure 1.**
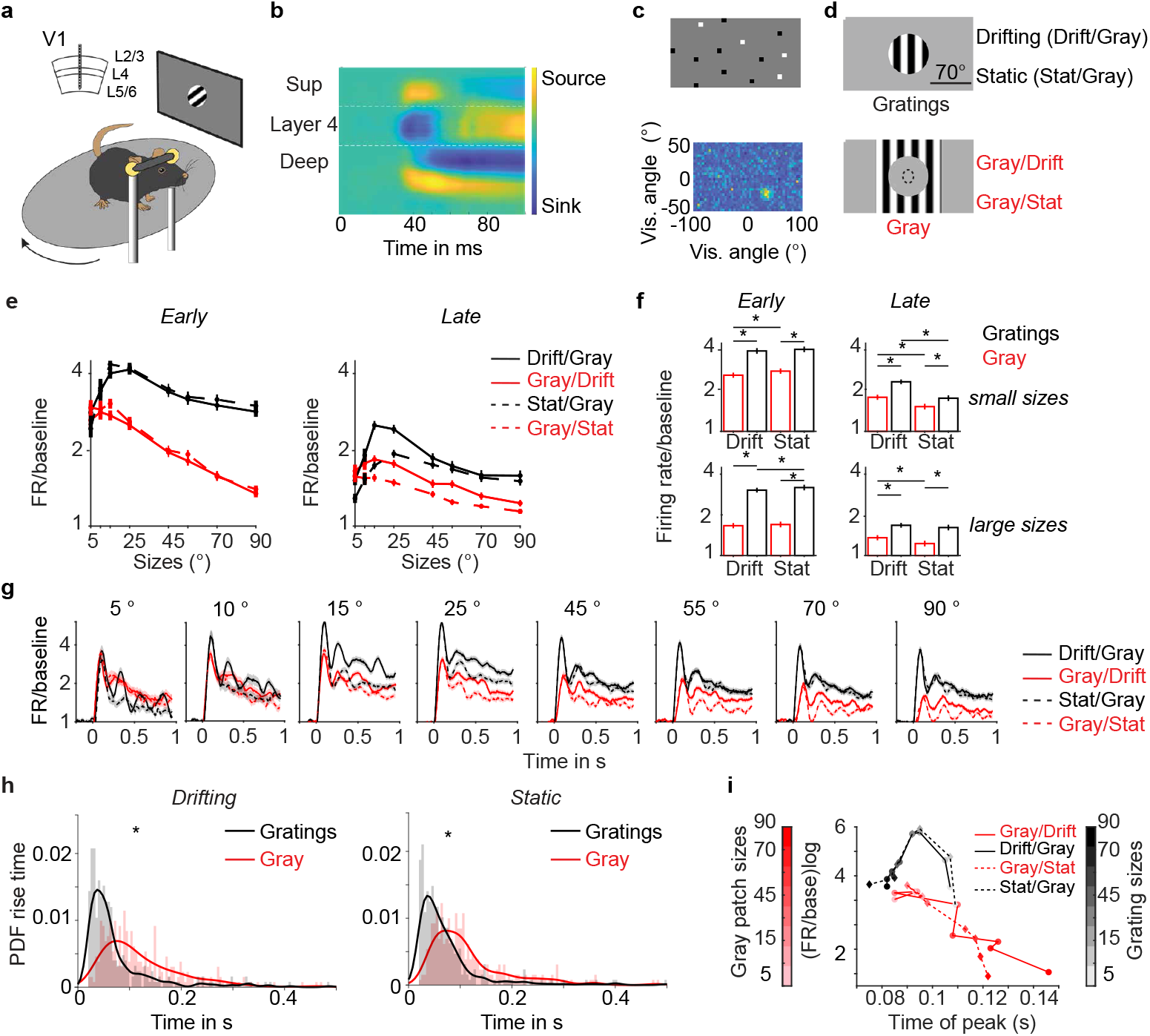
V1 responses to gray patches with gratings in the surround. a) Extracellular recordings across V1 layers in awake head-fixed mice on a running disk. b) Example session of current source density analysis to identify cortical layers. c) Sparse noise protocol (top) for RF (receptive field) mapping. Example RF for MUA (multi-unit activity). d) Main stimulus conditions: In the classical condition, gratings of different sizes were presented, either drifting (Drift) or stationary (Stat). In the gray condition, a gray patch was centered on the neuronal RF and had the same luminance as the background during the inter-trial interval (baseline). Hence, at stimulus onset, only the surround stimulus changes. Example of a grating (top) or gray center patch (bottom) of 70°. The dashed circle represents an RF of 20° diameter. e) Average firing rates of single units, normalized to baseline, shown in logarithmic scale. The left panel corresponds to the early (0.04 s to 0.15 s) and the right panel to the late stimulus period (0.2 s to 1 s after stimulus onset) (number of neurons *n* =335, 6 animals). f) Statistical analysis for all conditions from e). Sizes are separated into small *<* 45° and large ≥ 45° (*p-values *<* 0.01 comparing drifting vs. stationary per size, Wilcoxon signed-rank test). All conditions had values higher than the baseline (p-values < 0.01 Wilcoxon signed-rank test). g) Average spike density normalized to baseline. Solid lines represent drifting conditions, and dashed lines represent stationary conditions. The black line on top of each subplot represents the stimulus period. h) Histogram of rise times of neural responses for Gratings (black) or Gray (red) (sizes ≥ 45°). PDF is the probability density function. Solid lines are a Kernel smoothing function of the histogram (Wilcoxon signed-rank test, *p-value < 0.01). i) Each point represents the peak response time of the average spike density function as a function of response magnitude.

Throughout the paper, we use the following nomenclature: We will describe the configuration of a stimulus in the center, e.g., a circular gray patch superimposed onto a background consisting of a drifting grating, as “Gray/Drift”. Likewise, we shall refer to a grating superimposed onto a gray background as “Drift/Gray” or simply as “Drift”. In the first experimental paradigm, we presented both stationary and drifting grating stimuli. The stimuli were presented in four main conditions, namely “Drift/Gray”, “Stat/Gray”, “Gray/Drift” and “Gray/Stat” (Figure 1d). In the Drift/Gray and Stat/Gray conditions, gratings of different sizes were presented with direct visual stimulation of the neurons’ classical RFs with a grating stimulus. By contrast, in the Gray/Drift and Gray/Stat (patch) conditions, gray circular patches of different sizes were positioned at the center and a grating of a fixed size was presented in the surround (see Methods). The gray patches thus effectively occluded part of the grating stimulus and had the same intensity as the gray screen in the inter-trial interval. We separately analyzed the early (0.04 s to 0.15 s) and late (0.25 s to 1 s) neuronal responses and refer to these as the early stimulus period and late stimulus period.

In the Drift/Gray condition, neurons showed maximum firing rates for gratings of sizes around 15°-25° of diameter. Firing rates gradually decreased with the size of the drifting grating, i.e. surround suppression (Figure 1e-g, S1a-c). In the Gray/Drift condition, neuronal firing rates also reached a maximum value for gray patch sizes around 15° of diameter, with a decrease in firing for larger gray patch sizes (Figure 1e-g). We observed an increase in neuronal firing rates during the late stimulus period (but not during the early period) for a 15° diameter gray patch in the Gray/Drift condition, as compared to the response to the 90° grating stimulus in the Drift/Gray condition (Figure S2d). Keller et al. (2020) have described this firing increase in the Gray/Drift condition relative to the large drifting grating stimulus as an “inverse receptive field”.

Importantly, increased firing rates relative to baseline for gray patch sizes of 15°-25° diameter (Figure 1e-g) might potentially be explained by the presence of an edge in the neuronal receptive field. Because of variability in RF centers across neurons, the stimulus was not always exactly centered on the neuronal RF but could be 10° diameter away from the RF center. Consequently, for gray patch sizes of 15°-25° diameter, a small part of the grating stimulus may have been placed inside the classical RF in the Gray/Drift condition. This explanation does not apply to larger gray patch sizes, in which case we can be certain that the surround stimulus does not induce a direct bottom-up drive. In our experiment, we included gray patches with sizes up to 90°. Strikingly, we found that in the Gray/Drift condition, firing rates were increased relative to baseline (i.e. the fullfield gray screen) even for large gray patch sizes up to 90°, (Figure 1e-h). Thus, a stimulus presented in the distal surround induced a reliable increase in firing rates relative to baseline. We shall refer to this effect as the “surround-induced response”. Surround-induced responses were stronger in the early stimulus period (0.04 s to 0.15 s) than in the late stimulus period (from 0.2 s to 1 s) (Figure 1h).

We wondered if there would be a major difference in the magnitude of surround-induced responses between Gray/Drift and Gray/Stat conditions, considering that for a drifting grating, the visual system may infer that an object is moving behind the gray patch. However, surround-induced responses were found for both drifting and stationary grating stimuli presented in the distal surround. In the early stimulus period, in which surround-induced responses were the strongest, we did not observe a significant difference in the magnitude of surround-induced responses between Gray/Drift and Gray/Stat conditions. However, we did observe a stronger rate increase for the Gray/Drift condition as compared to the Gray/Stat condition in the late stimulus period (Figure 1e,f).

We further analyzed how the temporal structure of the surround-induced response differed from responses in the classical stimulus condition. To study the latencies of firing responses, we first quantified, for each neuron, the rise time of the neuronal response based on the spike-density function. We then compared response latencies between Gray/Drift (for a gray patch of 45° and larger) and Drift/Gray (for a drifting grating of 45° and larger). Response latencies were delayed for both the Gray/Drift and Gray/Stat conditions as compared to the Drift/Gray and Stat/Gray conditions (Figure 1h). For both the Gray/Drift and Gray/Stat condition, response latencies showed a systematic increase with the size of the gray patch, resulting in a negative correlation between the magnitude of the neuronal response and the response latency (Figure 1i).

Specifically, response latencies were comparable between the classic (i.e. Drift/Gray and Stat/Gray) and patch (i.e. Gray/Drift and Gray/Stat) conditions up to gray patch sizes of about 45° diameter (Figure 1h). For larger center stimuli, firing responses in the Gray/Drift and Gray/Stat conditions were delayed by about 50 ms compared to the classical Drift/Gray and Stat/Gray conditions (Figure 1h).

In addition, we recorded single LGN neurons using the same paradigm (Figure 2a,b). Similarly to V1, LGN neurons had a maximum response to drifting and static grating stimuli (i.e. Drift/Gray and Stat/Gray) for sizes around 15°-25° diameter. In contrast to V1 neurons, LGN neurons did not show an increase in firing rates relative to baseline for larger diameters of the gray-patch in the Gray/Drift and Gray/Stat conditions. Instead, during both the early and late stimulus period, LGN firing rates decreased below baseline levels, with maximum suppression for gray patches of 45° diameter (Figure 2c, Rank-Wilcoxon test, p<0.01). Even for gray patches of 25° diameter, the firing response of LGN neurons was weak and did not differ from baseline levels in the late stimulus period. These analyses indicate that the increase in V1 firing rates for gray-patch diameters of 25° and larger is not inherited from area LGN but depends on horizontal and top-down cortical feedback.

**Figure 2.**
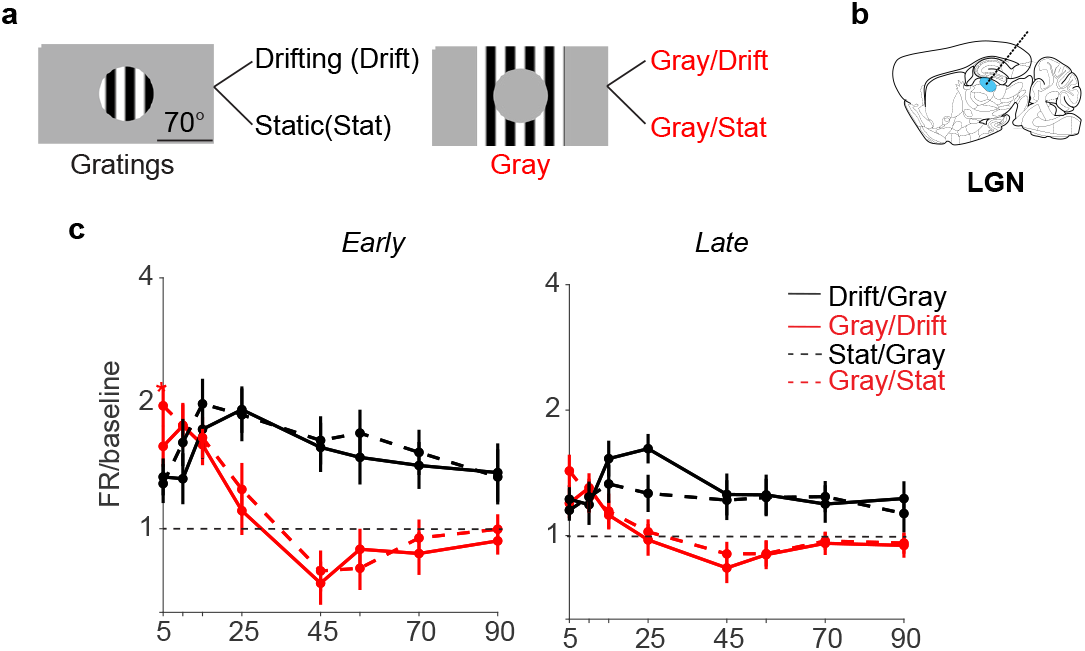
LGN responses to gray patches with a grating in the surround. a) Stimuli, as in Figure 1. b) Representative scheme of extracellular recordings in LGN. c) Average firing rates normalized to baseline (*n* =30, 2 animals).

### Responses induced by discontinuous surround stimuli

As described above, grating stimuli presented in the distal surround can increase V1 firing rates relative to baseline (i.e. surround-induced response). These grating stimuli were spatially coherent, i.e. they had a continuous spatial structure that was interrupted by the gray center patch. Such a continuous grating stimulus allows for prediction of the object occluded behind the gray center patch via interpolation. We therefore wondered to what extent the surround-induced response depends on the spatial continuity of the surround stimuli.

To investigate this, we disrupted the spatial continuity of the surround stimulus by dividing the surround stimulus into two separate gratings of orthogonal orientations (Figure 3a,b). These two gratings were placed next to each other, with the dividing line centered on the neuronal RF. In the drifting-grating condition, the stimuli moved in orthogonal directions. Gratings were presented in two conditions, either vertical and horizontal (0° and 90°) or diagonal (45° and 135°) (Figure 3a,b). In the vertical-horizontal condition, the two gratings were spatially discontinuous at the location of the patch, whereas in the diagonal condition, the gratings were spatially continuous (although the motion of the grating was non-coherent in both conditions). The grating stimuli could appear in four configurations: drifting/stationary (Drift/Stat) and continuous/discontinuous (Cont. or Disc.), i.e. Drift.Cont., Drift.Disc., Stat.Cont., and Stat.Disc. For the Gray/Grating condition, a rectangular gray patch was superimposed onto the grating stimuli. We varied the width of this rectangular gray patch up to a width of 90°.

**Figure 3.**
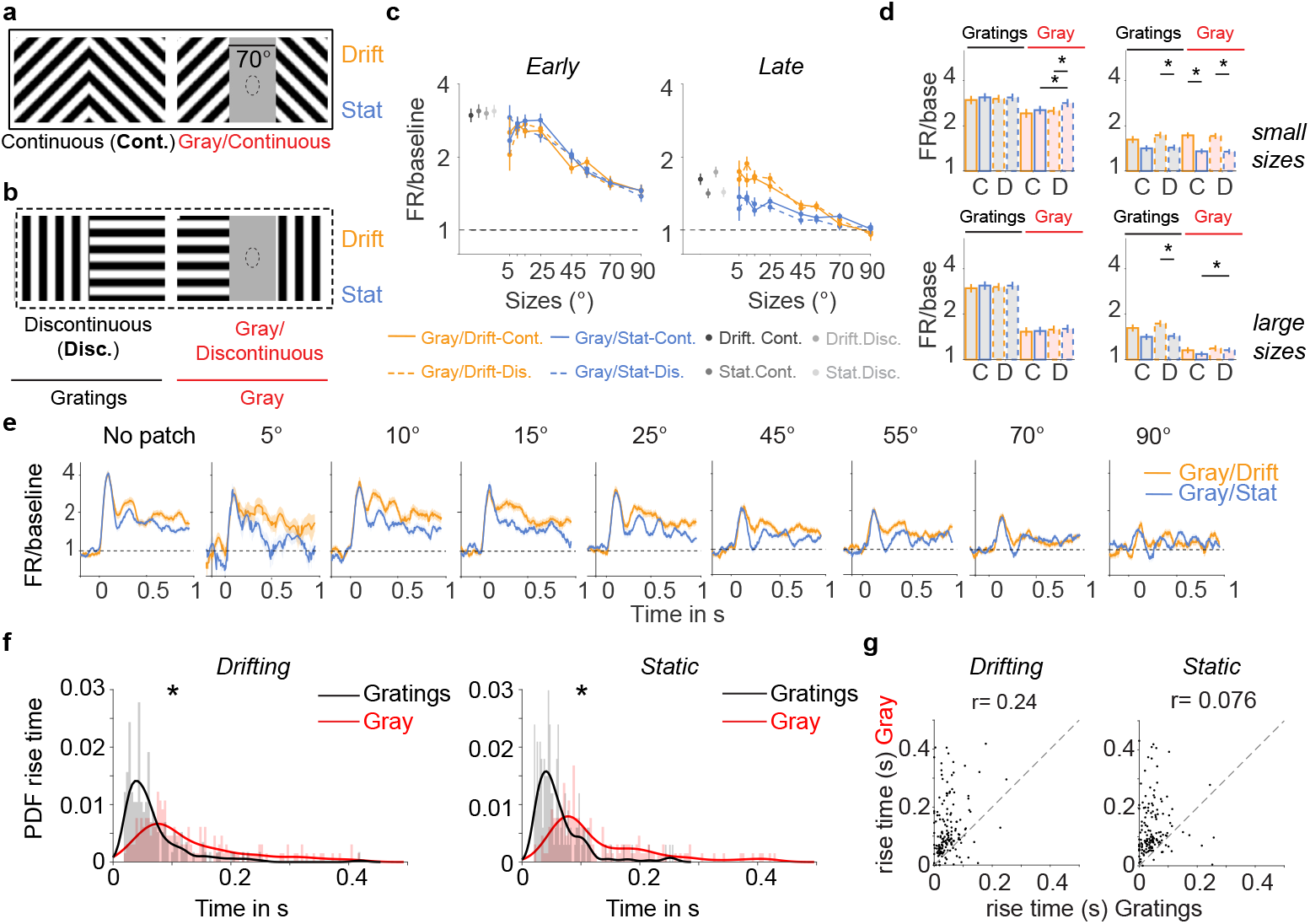
Neural responses to rectangular gray patches with orthogonal grating stimuli in the surround. a) Spatially continuous gratings and a gray rectangular patch covering the gratings (Gray/Cont). The stimulus could be presented as either drifting or stationary. b) Discontinuous gratings in the surround and discontinuous gratings covered by the gray patch (Gray/Disc). c) Population size tuning, shown as firing rates normalized to baseline during early (0.04 s to 0.15 s) and late (0.2 s to 1 s) stimulus periods. “Drift” and “Stat” refer to drifting and stationary conditions. The sizes correspond to the different dimensions of the rectangular patch. In this condition, the classical condition was presented only as full-field gratings. d) Statistical analysis of data in (d). Sizes were divided in small (*<* 45°) and large (≥ 45°) (*p-values *<* 0.01 Wilcoxon signed-rank test. *n* =132 single units in 5 animals). All conditions were significantly higher than baseline during early and late periods (p < 0.01, Wilcoxon signed-rank test). e) Average spike density function for different sizes of the rectangular gray patch. The solid line represents the stimulus period. f) Histogram of response latencies (rise time) for Gratings (black) and Gray (red) conditions. Latency was computed for sizes of ≥ 45°. The black and red lines are (kernel) smoothing estimates. Drifting and stationary conditions are pooled together. PDF corresponds to the probability density function (Wilcoxon signed-rank test, *p-value *<* 0.01). g) Scatter plot of rise time for the Gratings vs. Gray condition (sizes ≥ 45°, r-Pearson correlation value).

We found surround-induced responses when the surround stimulus was a grating with orthogonal orientations (Figure 3c-e), with delays in response latencies for both Gray/Drift and Gray/Stat conditions (Figure 3f,g). The surround-induced response was found both for drifting and stationary gratings in the surround and for both spatially continuous (i.e. diagonal) and discontinuous (vertical-horizontal) gratings in the surround (i.e. Gray/Drift-Cont, Gray/Drift-Disc., Gray/Stat-Cont., Gray/Stat-Disc.). In fact, surround-induced responses were greater for the discontinuous than continuous surround stimuli during the late stimulus period (comparing Gray/Stat-Drift. vs. Gray/Stat- Cont. in Figure 3d). Hence, the spatial continuity of the stimulus in the surround does not increase surround-induced responses, and surround-induced responses can be observed also in case of surround stimuli exhibiting incoherent motion.

The observation that the spatial continuity of the surround stimulus is not necessary to generate a surround-induced response predicts that surround-induced responses might also occur when the surround stimulus is a noisy texture. To test this, we presented pink-noise stimuli either in the classical condition (Noise) or with a circular gray patch centered on the neuronal RF (Gray/Noise); (Figure 4a). In the Gray/Noise condition, surround-induced responses were observed for gray patches up to 90° diameter (Figure 4b,c). Surround-induced responses were strongest in the early stimulus period, and their magnitude showed a negative dependence on the size of the gray patch (Figure 4c). We also observed a difference of ∼50 ms in response latency between the classical Noise and the Gray/Noise condition (Figure 4d,e).

**Figure 4.**
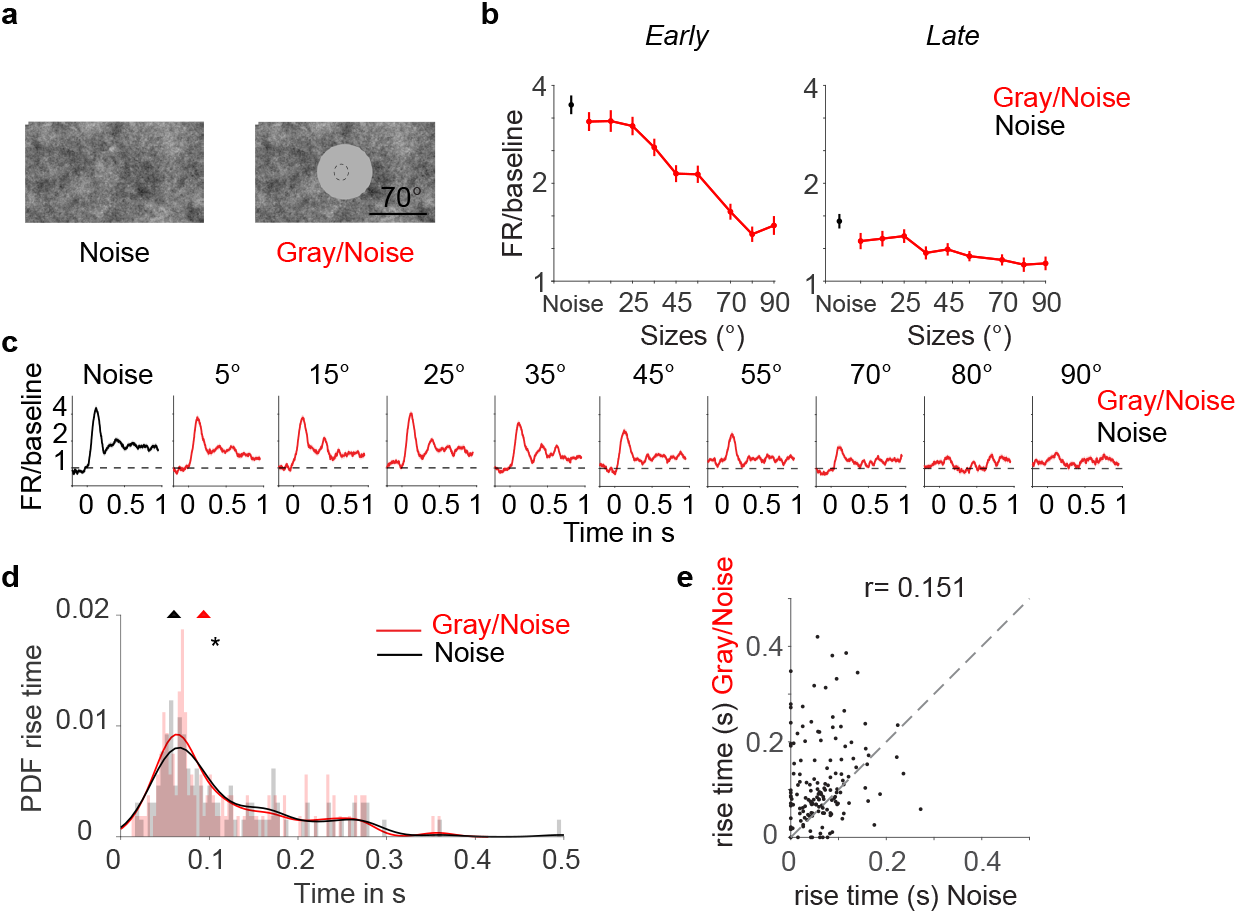
Neuronal responses to gray patch with pink noise background in the surround. a) Illustration of stimuli for 70° gray patch: Pink noise or a gray patch with pink noise background (Gray/Noise). b) Average firing rates (normalized to baseline) for different sizes of the gray patch in the gray condition. For comparison, we include the pink noise condition (PN; black dot). All sizes of the gray patch were significantly higher than the baseline, the comparison was performed in small (< 45 °) and large sizes (≥ 45°) during early and late stimulus periods (p < 0.01, Wilcoxon signed-rank test). c) Average (normalized to baseline) spike density function for different sizes of the gray patch. (*n* =139 units in 5 animals). d) Probability density function (PDF) of the rise time of the response. The line highlighted shows the Kernel smoothing function estimate from the PDF, and the triangles on top represent the median value for each population (Wilcoxon signed-rank test, *p-value *<* 0.01). e) Scatter plot of the rise time for the PN or Gray/Noise conditions (sizes ≥ 45°, r-Pearson correlation value).

### Uniform surface stimuli

If surround-induced responses comprise a representation of the gray surface centered on the RF, then one would expect that these responses can also be found when the gray patch is superimposed onto a uniform black or white background. Furthermore, it is plausible that responses to a gray surface stimulus on a white or black background show similarities to responses to a white or black surface stimulus on a gray background, both in terms of response latency and magnitude.

Similar to the experiments described above, we centered a gray patch on the neuronal RF. In this experiment, the background was either white or black (Gray/White and Gray/Black conditions) (Figure 5a,b). In the classical condition, we presented white or black patches of different sizes on a gray background, and we refer to these stimuli as white/black surface stimuli. In this case, the center of the stimulus was black or white, and the surround was gray (White/Gray; Black/Gray) (Figure 5a,b). We found that the onset of a white or black surround stimulus (Gray/White and Gray/Black) led to an increase in V1 firing rates above baseline levels. This firing increase was strongest for gray patches around 5°-15° diameter but was still significant for large gray patches (Figure 5c- f). V1 firing rates also increased for white or black surface stimuli centered on the neuronal RF (White/Gray and Black/Gray, Figure 5c-f). The differences in firing rates were relatively small between the Gray/White and White/Gray conditions and between the Gray/Black and Black/Gray conditions. In the late stimulus period, opposite patterns were found for white and black conditions: Firing rates were higher in the White/Gray than in the Gray/White condition (Figure 5e). However, firing responses were higher in the Gray/Black than the Black/Gray condition (Figure 5e). Hence, firing rates were generally higher in the condition with the brighter surface in the center. Furthermore, we found that latencies were not significantly different between conditions and peaked around 120 ms, similar to the latency observed for the Gray/Drift and Gray/Stat protocols (Figure 5g,h). Thus, firing responses to gray surface stimuli on a white/black background tend to have magnitudes and latencies that are similar to white/black surface stimuli on a gray background. This finding differs from the case of grating stimuli, where we found a strong latency difference and stronger response for grating stimuli on a gray background, as compared to gray patch stimuli on a grating background (see Figure S3 for a direct comparison).

**Figure 5.**
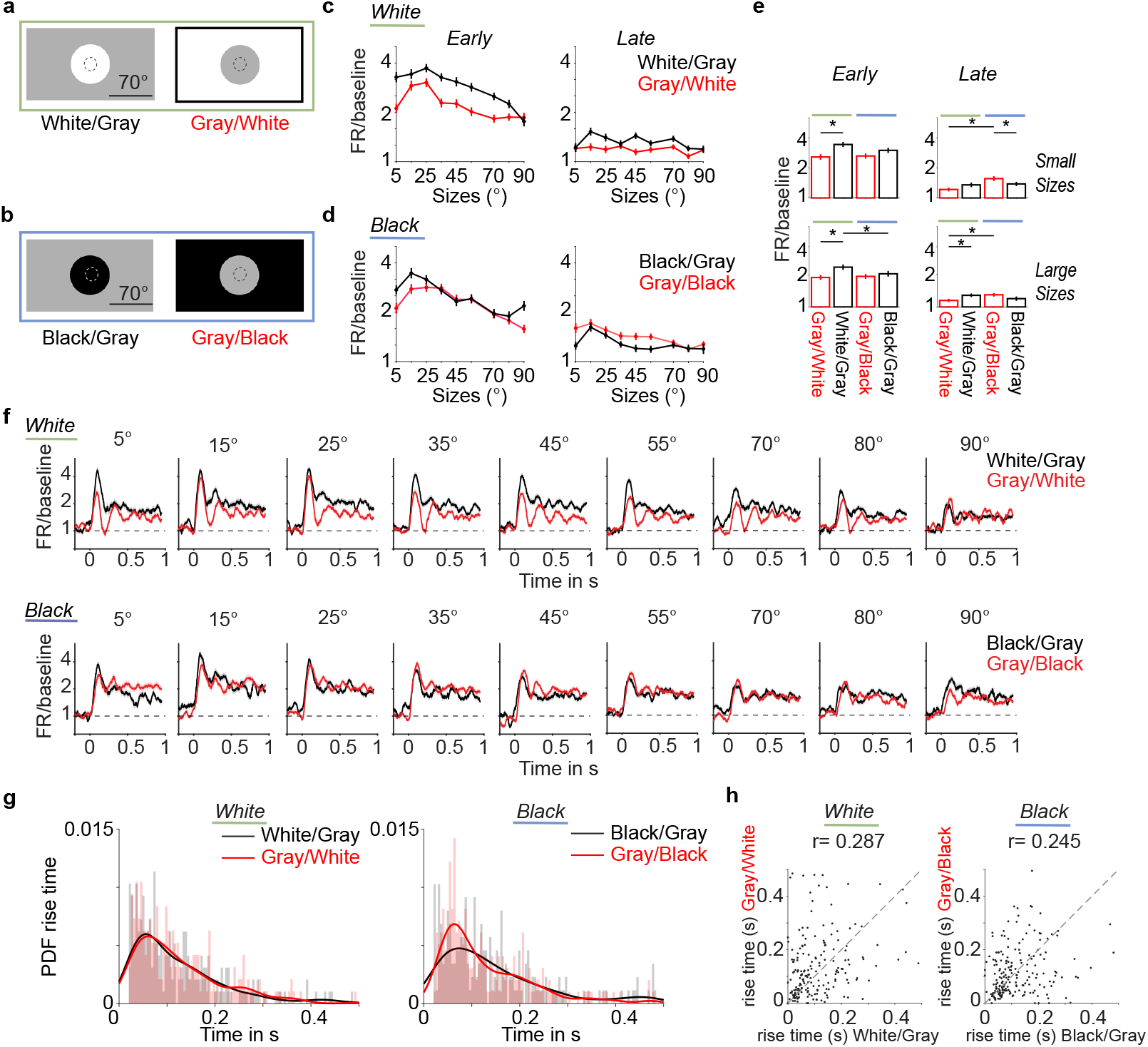
Neural activity for a gray patch with a black or white surround. a) White stimuli: white patches with a gray surround (White/Gray) or a gray patch with a white surround (Gray/White). b) Black stimuli: black center with a gray surround (Black/Gray) or a gray patch with a black surround (Gray/Black). c) Average firing rates normalized to baseline and in logarithmic scale for early (0.04 s to 0.15 s) and late (0.2 s to 1 s) stimulus period. d) Same as (c) for black stimuli shown in (b). e) Mean and SEM of neural responses for small and large patch sizes, separately for early and late stimulus periods. (*p-values *<* 0.01, Wilcoxon signed-rank test, *n* =170 units in 5 animals). All conditions were significantly higher than baseline. f) Average spike density for different stimulus sizes. Spike densities are normalized to baseline and shown in a logarithmic scale. g) Histogram of response latencies across neurons (response rise time). The line highlighted shows the estimated probability density (kernel smoothing). Response latencies were computed for patch sizes of 45° and larger. PDF corresponds to the probability density function. h) Scatter plot of the rise time for the classical and gray patch condition per unit (single units *n* =247, 5 animals, r-Pearson correlation value).

### Decoding of population firing rate vectors

Finally, we asked to what extent surround-induced responses carry information about the specific stimulus in the surround. To this end, we investigated differences in neuronal population vectors between different kinds of surround stimuli by computing the Euclidean distance between two firing rate vectors for all pairs of trials (Figure 6a-b). Based on these distance matrices, we computed low-dimensional embeddings via t-SNE (Figure 6c). In addition, we performed supervised classification via support vector machines (Figure 6d). We performed these analyses including 70° and 90° diameter gray patches or stimuli for the protocols with stationary and drifting gratings (Figure 1), the rectangular gray patch with orthogonal gratings (Figure 3, and the protocol with black/white backgrounds (Figure 5).

**Figure 6.**
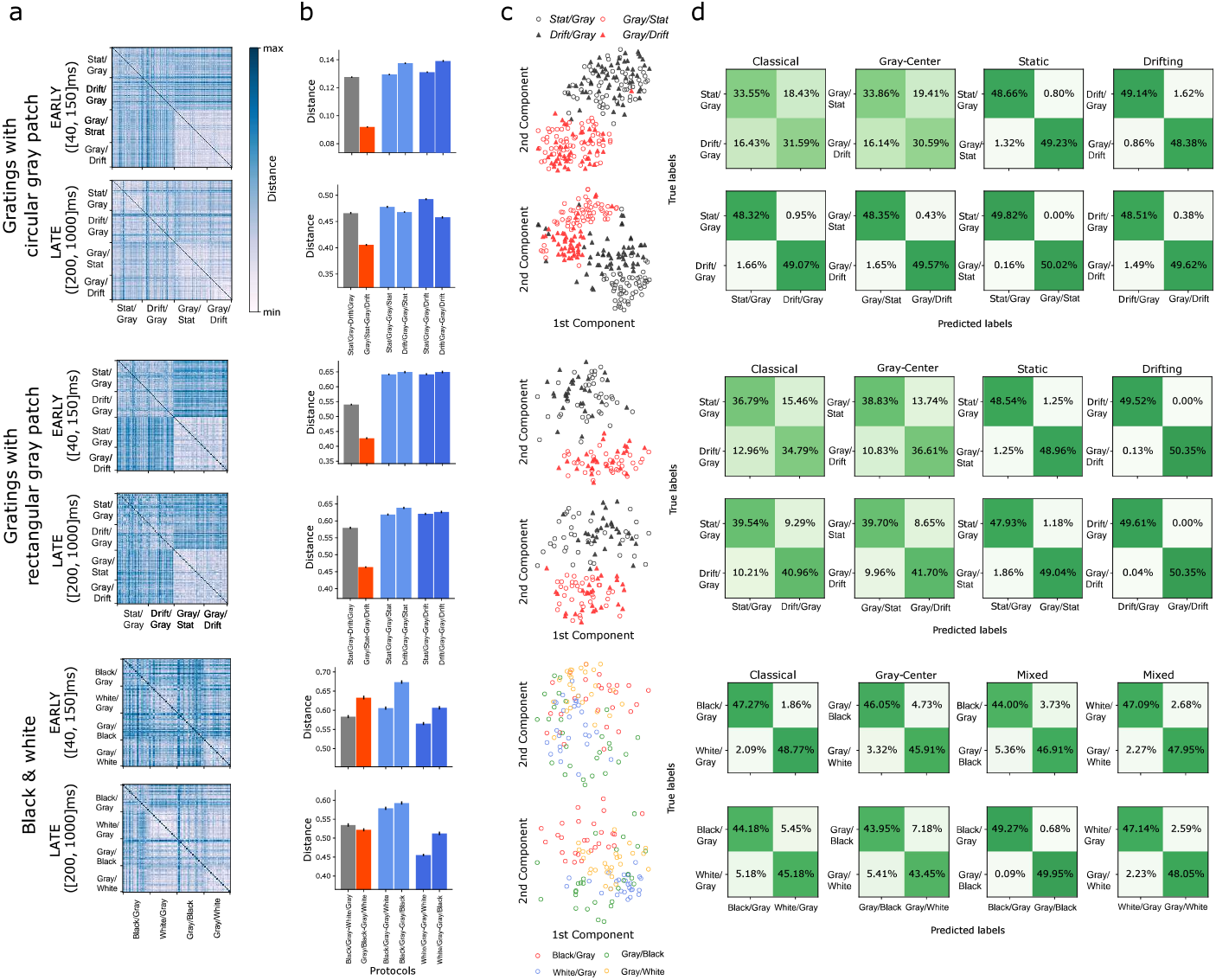
Population analyses to investigate stimulus specificity. a) Dissimilarity matrices of firing rate vectors across trials. The distance between firing rate vectors was computed using Euclidean distances. For visualization purposes, the diagonal shows the maximum value. b) Mean distances between protocols based on dissimilarity matrices. Black lines show the 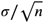 (i.e., SEM), where *n* is the number of samples (i.e., distances). Stat/Gray-Gray/Stat indicates the distance between stationary grating (classical) and gray-center/stationary-grating (patch) conditions. For gratings with a circular gray patch during early periods, only (Stat/Gray-Gray/Stat, Stat/Gray-Gray/Drift) were not distinguishable (p-val equals 0.396). For late periods, all the distances were statistically distinguishable. For Gratings with rectangular patch during early periods, (Stat/Gray-Gray/Stat, Drift/Gray-Gray/Stat), (Stat/Gray-Gray/Stat, Drift/Gray-Gray/Drift), and (Drift/Gray-Gray/Stat, Drift/Gray-Gray/Drift) were not significant distinguishable (p-values equal to 0.4585, 0.9402, 0.9478, respectively). For late periods, all the comparison yielded significance. Finally, for B&W, all the comparisons were statistically significant except for the distances between (Black/Gray-Gray/White, Gray/Black-Gray/White) (p-val = 0.0247) for early periods. For late periods, the comparisons (Black/Gray-Gray/Black, Black/Gray-White/Gray) and (Gray/Black-Gray/White, Gray/Black-White/Gray) were not significantly distinct (p-values 0.0338 and 0.0308, respectively). c) 2D t-SNE embedding based on dissimilarity matrices shown in a). d) Support Vector Classifier (SVC) based on matrices in a). Classification score across 20 repetitions. 40% of trials were used for training and 60% for testing.

Across protocols, we refer to the condition with the gray patch in the center as the “patch” condition, and the condition with the stimulus (grating, or black surface) centered on the RF as the “classical” condition. We first examined whether the patch and classical condition could be distinguished. In all three protocols, firing rate vectors formed distinct clusters for the patch condition and the classical condition, with classification performance above 90% for all stimulus conditions and early and late stimulus periods (Figure 6d). Next, we analyzed to what extent the surround stimulus in the patch condition could be decoded from the surround-induced response. For all protocols, the surround stimulus could be decoded with high accuracy during the early and late stimulus periods (Figure 6d). Decoding performances were comparable between the patch and classical conditions. That is, the surround-induced response contained about the same amount of information about the surround stimulus as the activity in the classical condition (i.e. when the same surround stimulus was presented) (Figure 6d).

Nevertheless, there were differences between the grating and rectangular protocol compared to the black and white protocol. The t-SNE and dissimilarity matrices showed two main clusters in the grating and rectangular protocol, one for the classical and one for the patch condition. This was reflected by the fact that the distance between stationary and drifting grating in the patch condition was substantially smaller than the distance between the other conditions. However, in the black-white protocol, we did not observe a distinct cluster for the gray patch condition, and the distance was not consistently lower than distances between the other conditions (Figure 6a-b). Finally, we examined if the surround-induced responses in the gray patch condition, for a given surround stimulus, tended to be similar to the responses in the classical condition for the same surround stimulus. This was generally not the case. However, the distance between a stationary grating in the classical and patch condition (Stat/Gray-Gray/Stat) was larger than the distance between a drifting grating in the classical and stationary grating in the patch condition (Drift/Gray-Gray/Stat), for example.

## Discussion

Recent studies suggest that V1 neurons can be driven by a surround stimulus when a gray patch covers the classical RF, an effect that likely depends on feedback (***Schnabel et al., 2018; Keller et al., 2020; Kirchberger et al., 2023***). The present study had two main objectives: First, to confirm that neural responses can indeed be driven by stimuli in the surround alone, by recording a large number of neurons electrophysiologically and using distal surround stimuli for which direct stimulation of the classic RF can be excluded. Our second aim was to systematically investigate the stimulus-dependence of surround-induced responses, thereby distinguishing different computational mechanisms and interpretations. In particular, previous work has proposed that surround-induced responses may result from predictive processing, either as a prediction of the occluded content or a prediction error (***Keller et al., 2020; Muckli et al., 2015***). A competing interpretation is that surround-induced responses reflect the representation of the uniform gray patch itself and relate to segmentation processes (***Zweig et al., 2015***).

We recorded V1 and LGN neurons using Neuropixels in awake mice and showed that V1 firing rates increase by presenting a grating stimulus in the background, while the RF is covered by a gray patch up to 90° of visual angle. Our findings suggest that the surround-induced responses are due to horizontal or top-down feedback because surround stimuli induced decreases in LGN firing rates, which had RF sizes comparable to V1. Furthermore, mice make infrequent and small eye movements, while our surround stimuli were up to 45° away from the RF center.

Increased firing at the gray center patch did not require spatial continuity or motion coherence of the surround stimulus and was generalized to noisy textures and uniform black or white surfaces in the surround. Responses to black/white surfaces (with a gray background) had a similar magnitude and response latency as responses to a gray patch with uniform black/white stimuli in the surround. V1 response latencies showed a systematic increase in the size of the gray center patch, similar to what we observed for black/white stimuli. Based on these findings, we suggest that increased V1 firing for a gray patch following the presentation of a distal surround stimulus primarily reflects the representation of the gray patch itself.

### Surround-induced responses in mice

Our results demonstrate that V1 neurons are driven (i.e. increased firing rates relative to baseline) by various kinds of distal surround stimuli. We refer to this effect as the “surround-induced response”. We further observed an increase in response latency with the increase in the size of the gray patch up to about 50 ms. In previous experiments in mice, ***Keller et al. (2020***) also presented drifting gratings masked by circular gray patches with sizes up to 90°. Some (their Figure 1) but not all (their Figure 4) of their figures showed increased ΔF/F activity above zero. We argue that surround-induced response should be distinguished from the concept of an inverse RF that was recently put forward in a mice study by ***Keller et al. (2020***). The surround-induced response reflects an increase in firing rates relative to baseline levels, while the inverse RF refers to an increase in firing for a full-field stimulus masked by a gray patch relative to the same full-field stimulus. Thus, the interpretation of the “inverse RF” is that V1 firing rates increase due to the omission of classic RF stimuli. However, it is unclear what mechanism underlies the inverse RF. ***Keller et al. (2020***) reported preferred inverse RF sizes of about 15°. Similarly, we found an increase in V1 firing rates as compared to the full-field grating when it was masked by gray circular patches of 15°. We observed this increase specifically in the late stimulus period and only for drifting, but not for static gratings (Figure S2). We further find that the neural response for stimuli around 15° to 25° does not have a delayed latency as compared to classic RF inputs (note that ***Keller et al. (2020***) did not analyze the dependence of response latencies on the gray patch size). Given that the strongest inverse response occurs for circular gray patches around 15° without a delayed response latency, it is possible that for such a stimulus there is some remaining bottom-up input into the classical RF. We noted that V1 RF sizes are approximately 15° large and that units are included in the analysis (also in ***Keller et al. (2020***)) that have a classic RF within 10° away of the stimulus center. Hence, some of the grating in the surround may still cover the classical RF, which is consistent with the finding that LGN neurons show increased firing relative to baseline in the gray-center/grating-surround condition for gray patches of 15°. We posit that when a small circular gray patch is superimposed onto a grating stimulus, there are two factors determining the neural response: (1) The gray patch changes the spatial frequency content of the classical RF input. Consequently, the bottom-up drive may decrease compared to a grating stimulus. (2) The mismatch between surround and classic RF input induced by the gray patch can increase the response strength. If the influence of the second factor exceeds the influence of the first factor, the overall neural response may increase compared to a full-field grating stimulus, giving rise to an “inverse RF”. As the second factor depends on feed- back, inverse RFs may be observed specifically in superficial layers (***Keller et al., 2020***).

However, it is unclear whether the phenomenon of surround-induced responses also occurs in primates. Several studies did not report surround-induced V1 firing responses for grating stimuli, neither in the anesthetized nor the awake monkey (***Gieselmann and Thiele, 2008; Slllito et al., 1995; Cavanaugh et al., 2002a***). By contrast, other studies did observe surround-induced V1 firing responses in primates. ***Rossi et al. (2001***) showed an increase in V1 firing rates in the awake monkey for an oriented textured surround with a gray patch of 4° centered around the neuronal RF. Similar to our study, this response was substantially weaker than the classical response and also delayed in time, and was induced by distal surround stimulation considering that macaque RFs are about 1° wide. ***Papale et al. (2023***) found increased V1 firing rates for natural scenes in macaques. In their study, the surround stimulation however occurred relatively close to the neuronal RFs. In humans, similar stimuli have been shown to increase V1 BOLD responses (***Muckli et al., 2015***).

It remains to be investigated what explains these discrepancies between non-human-primate studies. A possible explanation is that the studies reporting surround-induced responses in macaques used full-field stimulus in the surround (***Rossi et al., 2001; Papale et al., 2023***), similar to our study, while the other studies that did not report surround-induced responses used a smaller surrounding annulus (***Slllito et al., 1995; Cavanaugh et al., 2002a; Gieselmann and Thiele, 2008***). It is possible that many forms of surround stimulation induce subthreshold activity in V1 neurons, composed of a mixture of excitatory and inhibitory conductances, but that only a subset of stimuli induce suprathreshold activity. Indeed, studies of the cortical point-spread function demonstrate that a visual stimulus elicits a wave of activity propagating up to 10 times bigger than the size of the retinotopic start point (***Grinvald et al., 1994***). Moreover, studies in cats have shown that postsynaptic integration fields in V1 are up to five times larger than the integration fields of suprathreshold spiking activity (***Bringuier et al., 1999***). The latencies of the subthreshold potentials increased with the distance of the stimulus to the center of the integration field.

There may be structural differences between mice and monkeys that could entail differential contextual modulation: In contrast to macaque V1 (***Talluri et al., 2023***), neurons in mouse V1 can be strongly driven by many factors not related to visual stimulation (***Vinck et al., 2015; Stringer et al., 2019a***). Consequently, a modulatory or weak input caused by surround stimulation may lead to changes in suprathreshold activity in mice but not in monkey V1. Surround stimuli may have a different effect on inhibitory and excitatory neurons as compared to mice. One characteristic feature of primates is their high acuity vision as compared to rodents. Theoretical models of predictive and efficient coding entail that stimuli with high precision should induce stronger inhibitory feedback, whereas lower precision should lead to more pooling (***Huang and Paradiso, 2008; Coen-Cagli et al., 2012***). This is illustrated by the increased spatial summation for low-contrast stimuli compared to high-contrast stimuli (***Sceniak et al., 1999***). Therefore, there may be more spatial summation in mouse V1 than in monkey V1.

### Mechanisms

In the Introduction, we contrasted two interpretations of surround-induced responses:

1. Predictive processing accounts: A first possibility is that surround-induced response result from predictive processing. In this view, surround-induced responses could either represent a omission mismatch signal (i.e. prediction error) (***Keller et al., 2020; Rao and Ballard, 1999***), or a prediction of the content behind the mask (***Derrington, 1996; Komatsu, 2006; Muckli et al., 2015***). We argue that the prediction-error explanation may account for the inverse RF, but not for the surround-induced responses. For the surround-induced responses, there is no mismatch when there is a large gray patch centered around the neural RF, because the bottom-up input into the RF (i.e. homogeneous gray surface) is the same as the near (proximal) surround (i.e., also a homogeneous gray surface). In other words, for a 90° gray patch centered on a neuron’s classic RF of 15°, there is a 75° gray surround, such that the RF input should be entirely predicted from the surround. Furthermore, contrary to our observations, one would have expected that a mismatch response depends on the continuity of the surround stimuli, i.e. whether a consistent prediction based on the surround can be generated. Yet, we showed that the surround-induced response generalizes to moving and stationary stimuli, continuous and discontinuous stimuli, noisy textures, and uniform surfaces in the surround. We furthermore argue that our data does not support that surround-induced responses reflect a perceptual inference (prediction) of the stimulus content behind the gray patch (acting as an occluder): First, we found equally strong surround-induced responses when the surround stimulus was not spatially continuous. For such a stimulus, the rectangle is not perceptually interpreted as an occluder of a “hidden” object. Likewise, we did not find stronger surround-induced responses for moving stimuli in the early period, even though moving stimuli should facilitate the inference that there is an object behind the gray patch. Second, we found equally strong surround-induced responses when the gray patch appeared as a salient object over a non-salient uniform black or white background. In this case, it is unclear why surround-induced responses would represent the uniform background behind the salient object rather than the salient and directly visible object (the gray patch) itself.
2. Segmentation accounts: A second possibility is that the surround-induced responses reflect the representation of the uniform gray patch itself, and relate to segmentation processes A previous study in macaque V1 has shown that for uniform surfaces (e.g. a black patch on a gray background), neural firing increases at the center of the uniform surface with a delay (relative to stimulus onset) compared to the response at the edge (***Zweig et al., 2015; Peter et al., 2019***). This delay in V1 activity increased with the size of the black or white surface stimulus (***Zweig et al., 2015***). This effect was interpreted as the inference of the surface information itself. Importantly, this surface information may not be available from the direct feedforward input, considering that a uniform surface has zero power at all spatial frequencies (***Zweig et al., 2015***). Thus, according to ***Zweig et al. (2015***), the V1 representation of the center of a uniform surface stimulus derives from neural responses at the edge of the surface.

We argue that the surround-induced response with a gray patch in the RF has a similar mechanistic origin and may reflects a representation of the gray patch itself. That is, presenting a distal surround stimulus activates neurons around the edge of the gray patch, which then leads to a transient and delayed increase in V1 firing (i.e. a surround-induced response) at the center of the gray patch. This interpretation is compatible with several observations: First, we showed that the response magnitude and latency of the surround-induced response were very similar to the neural response when a black or white patch was presented on a gray background. In fact, in the late stimulus period, surround-induced responses (i.e. with a gray patch) were stronger than responses to a black patch on a gray background. Furthermore, we did not observe that the population vectors for the gray patch on a black or white background formed a separate cluster (in the t-SNE embedding) as compared to a black or white patch on a gray background. Second, similar to ***Zweig et al. (2015***), we observed a systematic increase in the latency of surround-induced responses as a function of the surface (patch) size.

A closely related explanation is that the surround-induced responses represent a figure-ground effect (***Self et al., 2013; Schnabel et al., 2018; Kirchberger et al., 2023***) as suggested in mice and macaque studies. In this interpretation, the surround-induced response occurs because the gray patch appears as the figure (i.e. the foreground) on a background, and thus draws bottom-up attention (i.e. is salient). While figure-ground modulation may have contributed to the increase in V1 firing, we note that figure-ground modulation assumes that there is some representation of the gray patch to begin with, begging the question of how this representation emerges. Following (***Zweig et al., 2015***), we argue that the representation of a uniform surface stimulus, with information traveling from the edge to the center, forms a mechanism through which the surface is seen as an object, leading to perceptual grouping and image segmentation. These signals can then be further boosted when the surface appears as a figure on a background (i.e. figure-ground), however, they may also occur when e.g. the patch is large and flanked by two salient stimuli, as observed here.

It is possible however that V1 representations are mixed and reflect both segmentation and predictive processes. In this way, a single V1 vector could contain information both about the stimulus itself (i.e. the gray patch) but also about the spatial context in which it is embedded. That is, surround-induced responses may not merely encode the surface information of the gray patch, but could in addition encode information about the properties of the distal surround stimulus. Such a scenario would be consistent with the finding that human fMRI activity contains information about the predicted content behind the occluder (***Muckli et al., 2015***). In our study, we did observe that the surround-induced response had some degree of stimulus-specificity: We showed that it was possible to decode with high accuracy if the surround stimulus was drifting or stationary. Likewise, it was possible to decode if the surround stimulus was black or white. It is possible however that e.g. the difference between a stationary and drifting grating reflects the strength of the surround input, with less adaptation for drifting surround stimuli. Thus, more work is required to investigate this stimulus-specificity and distinguish e.g. adaptation from predictive processing accounts.

In sum, the most consistent explanation for our empirical observations of increased V1 firing due to a distal surround stimulus is that the distal surround stimulus evokes a representation of the gray center patch covering the classical RF, which can contribute to segmentation processes.

## Methods and Materials

### Materials availability

Further information and requests for resources should be directed to Martin Vinck (martin.vinck@esi- frankfurt.de).

### Data and code availability

The open-source MATLAB toolbox Fieldtrip (***Oostenveld et al., 2011***) was used for data analysis. Data and custom MATLAB scripts are available upon request from Martin Vinck (martin.vinck@esi- frankfurt.de) or Nisa Cuevas. For the population analysis, we used Scikit-Learn 0.22.1, Numpy 1.18.1 and Numba 0.51.2 for data cleaning and multi-CPU processing, SciPy 1.5.4 for statistics, and Matplotlib 3.1.3 for visualizations.

### Animals

The experiments were conducted in compliance with the European Communities Council Directive 2010/63/EC and the German Law for Protection of Animals, ensuring that all procedures were ethical and humane. All procedures were approved by local authorities, following appropriate ethics review. We included female and male mice (C57BL/6), a total of six animals for V1 recordings between three and eight months old. In two of those animals, we recorded simultaneously from LGN and V1. In one of the animals, we only recorded the gratings protocol, hence, that protocol has 6 animals, and the other protocols have 5 animals. Mice were maintained on an inverted 12/12 h light cycle, and recordings were performed during their dark (awake) cycle.

### Head Post Implantation Surgery

One day before the surgery, we handled the mice to reduce stress on the surgery day. We administered an analgesic (Metamizole, 200 mg/kg, sc) and an antibiotic (Enrofloxacin, 10 mg/kg, sc, Bayer, Leverkusen, Germany) and waited for 30 minutes. Anesthesia was then induced by placing the mice in an isoflurane-filled chamber (3% in oxygen, CP-Pharma, Burgdorf, Germany) and maintained throughout the surgery with isoflurane (0.8-1.5% in oxygen). We regulated the animal’s body temperature by using a heating pad, previously set to the body temperature. We constantly applied eye ointment (Bepanthen, Bayer, Leverkusen, Germany) to prevent eye dryness. Before making an incision, the skin was disinfected three times with Chlorexidine, followed by ethanol each time. After exposing the skull, we cleaned it with 3% peroxide three times, followed by iodine each time. The animal was positioned on a stereotaxic frame (David Kopf Instruments, Tujunga, California, USA). The skull was then aligned, and we measured the coordinates for V1 bilaterally, utilizing the transverse sinus as a reference point as previously described (***Wang et al., 2011***) (V1, AP: 1.1 mm anterior to the anterior border of the transverse sinus, ML: 2.0-2.5 mm) and marked the coordinates for V1. We positioned a screw in the frontal part of the skull to stabilize the implant. A custom-made titanium head-post was placed at the level of bregma, securing it with dental cement (Super-Bond C & B, Sun Medical, Shiga, Japan). The area designated as V1 was covered using cyanoacrylate glue (Insta-Cure, Bob Smith Industries Inc, Atascadero, CA USA). We closely monitored the animal’s recovery for 3-5 days, administering antibiotics for two consecutive days and providing metamizole in drinking water. We acclimated the animals to the running disk over five days. On the first day, we placed the mice on the disk for 5 minutes in complete darkness. We gradually increased the duration of exposure over the following days.

### Extracellular Recordings

On the day of the recording session, we performed a circular craniotomy of approximately 0.8 mm-1 mm diameter on V1 while the animals were under anesthesia (Isoflurane). We administered dexamethasone and metamizole thirty minutes before the procedure. We covered the craniotomy with Kwik-Cast (World Precision Instruments, Sarasota, USA) and inserted two pins into the cerebellum for grounding. We waited for at least 2 hours before the recording session. For the recording sessions, awake animals were head-fixed and placed on a running disk. We used Neuropixel probes, the probe was inserted around 1100-1300 µm depth with a 15° angle and recorded simultaneously from ∼150 channels, for LGN recordings we simultaneously recorded 384 channels. For each animal, we recorded around 2-3 sessions from each hemisphere, and we recorded from both hemispheres. For histological confirmation, we coated the probe in DiD (Invitrogen, 1 mg/mL) before the recordings to track the location of the probe. We isolated single units with Kilosort 2.5 (***Steinmetz et al., 2021***) and manually curated them with Phy2 (***Rossant et al., 2021***). We included only single units with a maximum contamination of 10 percent.

### Visual stimuli

The experiment was run on Windows 10 and stimuli were presented on an Asus PG279Q monitor set at 144 Hz refresh rate, racing mode, contrast 50% and brightness 25%. We employed Psychtoolbox-3 (***Brainard, 1997***) to create the stimuli presented. Throughout the study, we consistently placed the screen at a 30° angle of the eye contralateral to the recording hemisphere at a distance of 15 cm. For all protocols, the stimulus duration was 1 s, followed by an inter-trial interval of 1.3 s unless specified.

#### Sparse Noise and Receptive Field Mapping

We employed a locally sparse noise protocol to find the center of the RFs, modified from Allen Brain (see https://observatory.brain-map.org). The protocol consisted of black and white squares of 4.65 degrees, arranged in a 23×42 array. The stimulus was presented for 0.25 s, during which black and white squares were randomly positioned on a gray background. The total session duration was 15 minutes. We computed the response for each position separately, by averaging the response across all trials where a square was presented at a given position. A heatmap of the response was computed. This heatmap was then smoothed, and we calculated the location of the peak response. From the heatmap we calculated the centroid of the response using the function regionprops.m that finds unique objects, we then selected the biggest area detected. Using the centroids provided as output. We then fitted an ellipse centered on this peak response location to the smoothed heatmap using the MATLAB function ellipse.m. To center the visual stimuli during the recording session, we averaged the multiunit activity across the responsive channels and positioned the stimulus at the center of the ellipse fit to the MUA response averaged across channels. For all the following analyses based on the neuronal response to visual stimuli, we performed RF mapping using single-unit responses. During each trial, we collected responses to black and white squares presented in random positions on the screen and gray regions in the surround area not covered by a black/white square. We used a permutation test to compare the neuron’s responses to black and white squares inside the RF to the condition where there was no square in the RF (i.e. the RF was covered by the gray background). We included RFs of single units that met the following criteria: z-score of the response *>* 4, a permutation test p-value *<* 0.03, and an RF diameter within the range of 10° to 30°. We only included units in which the center of the RF was *<* 10° of visual angle from the center of the stimulus. As the locations of LGN RFs change across the dorsoventral positions, for LGN recordings, we averaged only channels with RFs close by and centered the gray patch’s position there. In each LGN recording session, we changed the locations of the center of the stimulus to two to three different positions.

#### Sinusoidal gratings

We presented drifting (2 cycles/sec) and static sinusoidal gratings, with a spatial frequency of 0.04 cycles per degree, with randomized orientations (0°, 45°, 90°, 135°, 180°, 225°) and sizes (5°, 10°, 15°, 25°, 45°, 55°, 70°, and 90°), equally balanced between gratings and gray patches over gratings. All stimuli were displayed in full contrast with a gray background. The patch had the same gray value as the one presented during the inter-stimulus interval. For the patch condition, we displayed gratings covering half the size of the x-axis of the screen. We presented only half of the x-axis due to the large size of our monitor, in order to avoid over-stimulation of the animals with very large grating stimuli. We presented 10-20 repetitions of each condition (2 motion conditions, 6 orientations, 8 sizes, 2 conditions of the patch, with or without a patch, in total 192 conditions per session). Luminance of all the stimuli were measured with Flame UV-VIS Miniature Spectrometer sensor placed at the center of the visual stimulus patch. Luminance intensities were constant across all stimulus conditions (100 lumen *cd*/*m*^2^).

#### Orthogonal gratings with elongated patch

We presented gratings with orthogonal orientations in each half size of the screen (Figure 3a-b). The drifting (0.04 cycles per degree) or static gratings were in randomized orientations (0°, 45°, 90°, 135°, 180°). We randomized conditions with full-field gratings without a patch (0°) or with a rectangular gray patch with different sizes of diameter (5°, 10°,15°,25°,45°,55°,70°, 90°). The sizes represent the varying dimensions of the rectangular patch. In this condition, the classical condition was shown only as full-field gratings, which is depicted in the plot as size 0, indicating no rectangular patch was present. For continuous gratings, the direction of the gratings on each side of the screen allowed for the completion of a pattern (one-half of the screen with 45° gratings and the other half of the screen with 135° gratings). Opposite, for the non-continuous condition, the orientations of gratings in each half of the screen did not allow pattern completion as one side was horizontal and the other side was vertical (0° *vs*. 90°). We presented 10-15 repetitions of each condition (8 sizes, 5 orientations and 2 stimuli conditions only gratings or gratings with patch, in total 80 conditions per session).

#### Pink Noise

We randomly presented one of two different pink noise images, together with a gray patch of different diameter sizes (0°, 5°, 15°, 25°, 35°, 45°, 55°, 70°, 80°, and 90°). We used two (high/low) contrast values of the pink noise, randomized, and each size of the patch was presented in 10-20 repetitions per session.

#### Black and white stimuli with patch

We showed 2 sets of stimuli: (1) White patches (centered on the RF) with a gray surround (WcGs) or a gray patch with a white surround (GcWs). (2) A black patch with a gray surround (BcGs) or a gray patch with a black surround (GcBs). The diameter size of the center patch was randomized (5°, 15°, 25°, 35°, 45°, 55°, 70°, 80°, and 90°), as well as the color (black or white) of the patch or the background. We presented around 10-15 repetitions per condition (9 sizes, and 4 conditions of the patch, either gray patch with white/black background or black/white patch with gray surround).

### Assignment of cortical layers in V1

The assignment of superficial, L4, and deep cortical layers was based on the current source density (CSD) of the average LFP signal during whole screen flash stimulation. The protocol consisted of a 100 ms long white screen period with a 2 s gray screen for the inter-stimulus period. To increase the spatial sampling rate, we interpolated the LFP traces with an interpolation factor of 4. CSD analysis was computed by taking the second discrete spatial derivative across the different electrode recording sites. The step size of the discrete spatial derivative was 200 µm. Single units were assigned to a cortical layer based on the location of the channel with the highest amplitude during a spike.

### Assignment of units in LGN

A flash stimulus was employed to confirm the locations of LGN at the beginning of the recording sessions, similar to our previous work in which we recorded from LGN and V1 simultaneously (***Schneider et al., 2023***). This stimulus consisted of a 100 ms white screen and a 2 s gray screen as the inter-stimulus interval, designed to identify visually responsive areas. The responses of multiunit activity (MUA) to the flash stimulus were extracted and a CSD analysis was then performed on the MUA, sampling every two channels. The resulting CSD profiles were plotted to identify channels corresponding to the LGN. During LGN recordings, simultaneous recordings were made from V1, revealing visually responsive areas interspersed with non-responsive channels. For LGN recordings, only the protocol with gratings was presented.

### Inclusion criteria

We included the following criteria in the spike-sorted units: 1) The ZETA-test (***Montijn et al., 2021***) was applied to the period around the onset of the classical gratings (0 ms, 250 ms) to test which neurons showed significantly modulated spiking activity (p-value<0.05 and zeta responsiveness *>* 2). 2) V1 units: assignment of the layer with CSD analysis. 3) Units that met the selection criteria of a good RF and Euclidean distance from the center of the RF to the center of stimulus had to lie within *<* 10 of visual angle. 4) Modulation of response to each protocol (gratings, black/white, pink Noise, and rectangular patch). We included units that were positively modulated for the classical condition of each protocol. The modulation response was calculated as the average firing rate during the stimulus presentation (30 to 250 ms) subtracting the average response from baseline (−250 to -30 ms) and dividing by the average response from baseline (i.e., (*F R*_*stim*_ *− F R*_*base*_)/*F R*_*base*_). For each stimulus protocol, we used the same units to compare different stimulus conditions (i.e., drifting vs. static).

### Statistical Analysis

We obtained the average firing rate from 0.04 s to 0.15 s for the early period and 0.2 s to 1s for the late period in the size-tuning plots. For all analyses, we normalized the responses per unit to the baseline, calculated the logarithm of the normalized responses, and presented the mean and standard error of the mean (SEM). For the spike density function for the different conditions, we used a time window of the Gaussian smoothing kernel from -.05 s to .05 s, with a standard deviation of 0.0125 s. The spike density functions of every unit were also normalized to the baseline, and we obtained the logarithmic values and presented them as mean responses and SEM. We defined the rise time (latency of responses) as the time in which the response of every unit (baseline subtracted) crossed a threshold 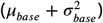 up to 0.5s. We plotted the population density function (PDF) of the rise times for diameter sizes of the patch or the gratings ≥ 45°. We included values *>* 0.02 s and obtained the kernel density function of the PDF. At the end, we calculated the Pearson’s correlation coefficient to correlate the rise time values of different conditions of visual stimuli. For all the statistical analyses, we calculated the Wilcoxon signed-rank test.

### Population Analysis

In total, the dataset yielded population spiking patterns that consisted of *N* = 344 neurons, which were pooled across multiple sessions as in previous studies (***Kheradpezhouh et al., 2020; Deitch et al., 2021; Sotomayor-Gómez et al., 2023***). For the population analyses, we analyzed the conditions in which the gray patch sizes were 70 ° and 90 °.

We calculated firing rate vectors for each analysis period by dividing the spike count per neuron by a window length *T*. From each dissimilarity matrix, we computed a 2D representation of epochs using the t-Distributed Stochastic Neighbor Embedding (t-SNE) manifold algorithm. For the t-SNE visualization, we used a perplexity value of 20 for the Gratings with circular and rectangular occluders, and 100 for the black and white condition. Although t-SNE is commonly employed for clustering tasks, our primary use of t-SNE was for visualization purposes. This allowed us to effectively represent the overall structure of the dissimilarity-based embeddings. Notably, changing the perplexity value would not influence the core analytical steps of our study, as t-SNE’s role was strictly to aid in visualizing group separation, not to impact the dissimilarity matrices or classification results.

We trained a C-Support Vector Classifier (C-SVC) based on dissimilarity matrices, which were calculated using Euclidean distance between firing rate vectors for all pairs of trials. The classifier was trained using 40% of the trials for training and 60% for testing. To ensure robust performance, we shuffled the trials for training and testing 20 times, repeating this procedure to account for potential variability. The classifier was binary, distinguishing between two classes (e.g., Dr vs St), and the classification score corresponds to the average accuracy across these 20 iterations.

### Statistical significance

We compared the population of distances, shown in Figure 6, using a two-sided Wilcoxon test. We computed the p-value for statistical comparison for gratings, gratings with the rectangular patch, and black and white stimuli. We consider *p*-*value <* 0.01 as a threshold for statistical significance.

### Face movement analysis

The mouse videos were reduced in dimensionality using SVD method as described in (***Stringer et al., 2019b***)

#### Video Acquisition

Infrared videos were acquired during the recording using either DALSA Genie Nano-M1450 or DALSA Genie Nano-M1280 GigE camera with zoom lens and an infrared filter (720 nm, Edmond optics R-72 cutoff). Outputted Camera exposure of each frame were used to synchronize video timing with the recording setup.

#### Pre-processing

Videos were cropped around the face of the animal and resized by 0.5. The absolute motion energy was computed as the absolute value of the difference between consecutive frames.

#### SVD

The absolute motion energy was subtracted by the average motion across all frames. Next, we computed the singular value decomposition (SVD) on the motion energy movie. The movie PCs were computed from svd output, and the top 4 PCs were used. We synchronized the videos to the behavioral paradigm and we calculated the motion signal of the trials with the gray patch, trials with gratings and intertrial intervals in which we presented a gray screen. The activity was normalized as

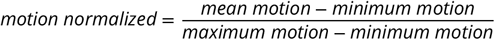

We considered trials with movement to trials that crossed a threshold calculated with the motion signal from the baseline trials (gray screen presented) as: *threshold* = *mean motion* + *Std Dev motion*^2^.

## Acknowledgments

Conceptualization: NC, MV. Experiments: NC, AT, AB. Data analysis: NC, BSG. Supervision: MV. Writing of main draft: NC, BSG, and MV, with comments from other authors. This project was financed by the BMF (Bundesministerium fuer Bildung und Forschung), Computational Life Sciences, project BINDA (031L0167); an ERC starting grant (850861) SPATEMP; DFG VI Grants (908/5-1 and 908/7-1, 505660261, 520285844); an NWO VIDI Grant; the Dutch Brain Interface Initiative (DBI2).

---

**Figure S1.**
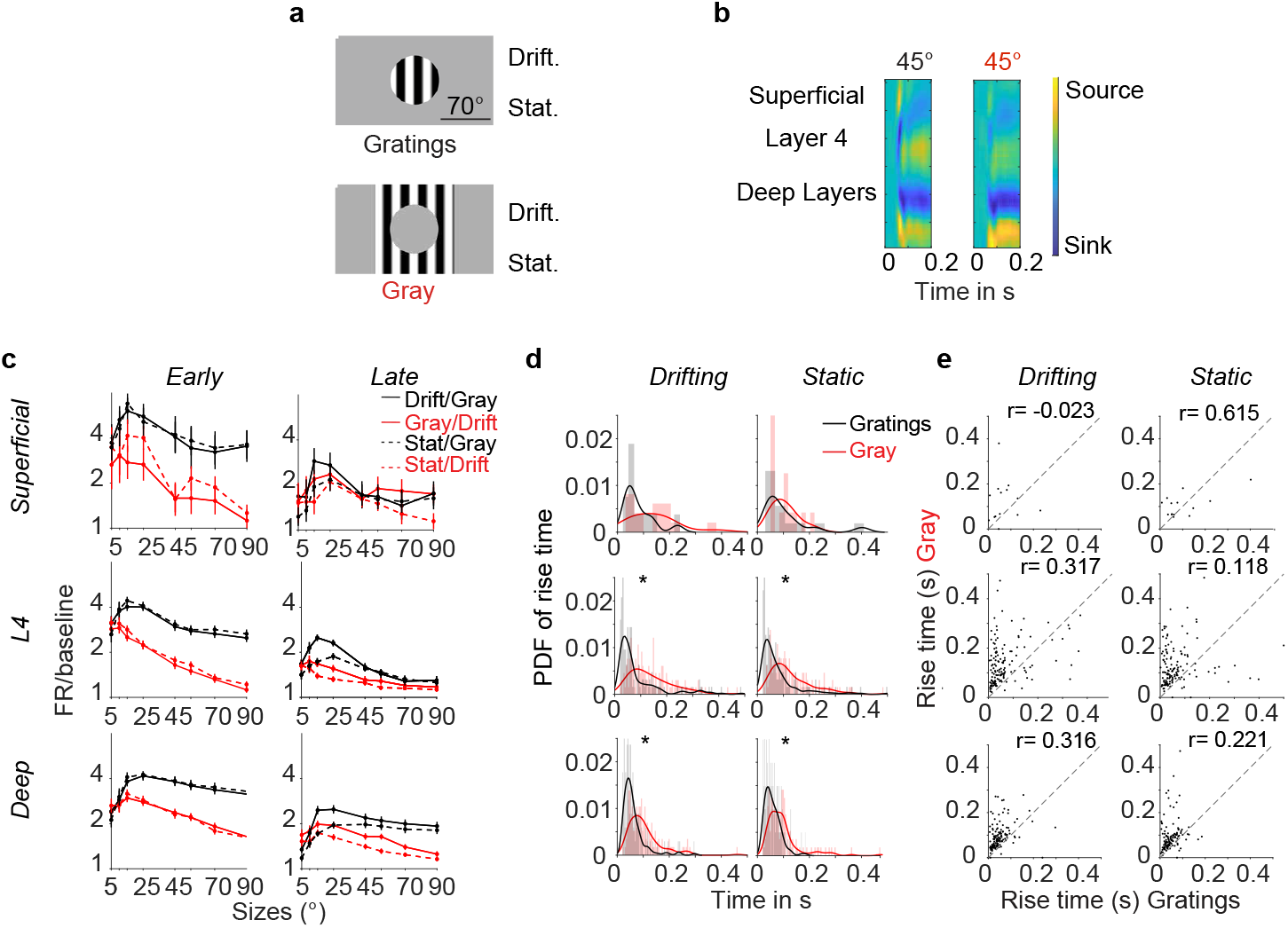
Firing rate across V1 Layers to gratings in the far surround a) Stimuli presented as in Figure 1). Drifting and stationary gratings and gratings covered with a gray patch. b) CSD analysis from the first 200 ms in response to 45° gratings (left) and gratings covered by a 45° gray patch (right). c) Population size tuning per layer. Firing rate during the early and late period of stimulation, every unit is normalized to baseline (Superficial units *n* = 22, L4 units *n* = 213 and deep layer units *n* = 208, Wilcoxon signed ranked test *p <* 0.01). d) Probability density function of the rise time per unit separated into layers for drifting and stationary gratings. From top to bottom superficial units, layer 4 units, and deep units (*p-values *<* 0.01 Wilcoxon signed ranked test *p <* 0.01). e) Scatter plot of the rise time for Gratings or Gray separated by layers (sizes ≥ 45°, r-Pearson correlation value).

**Figure S2.**
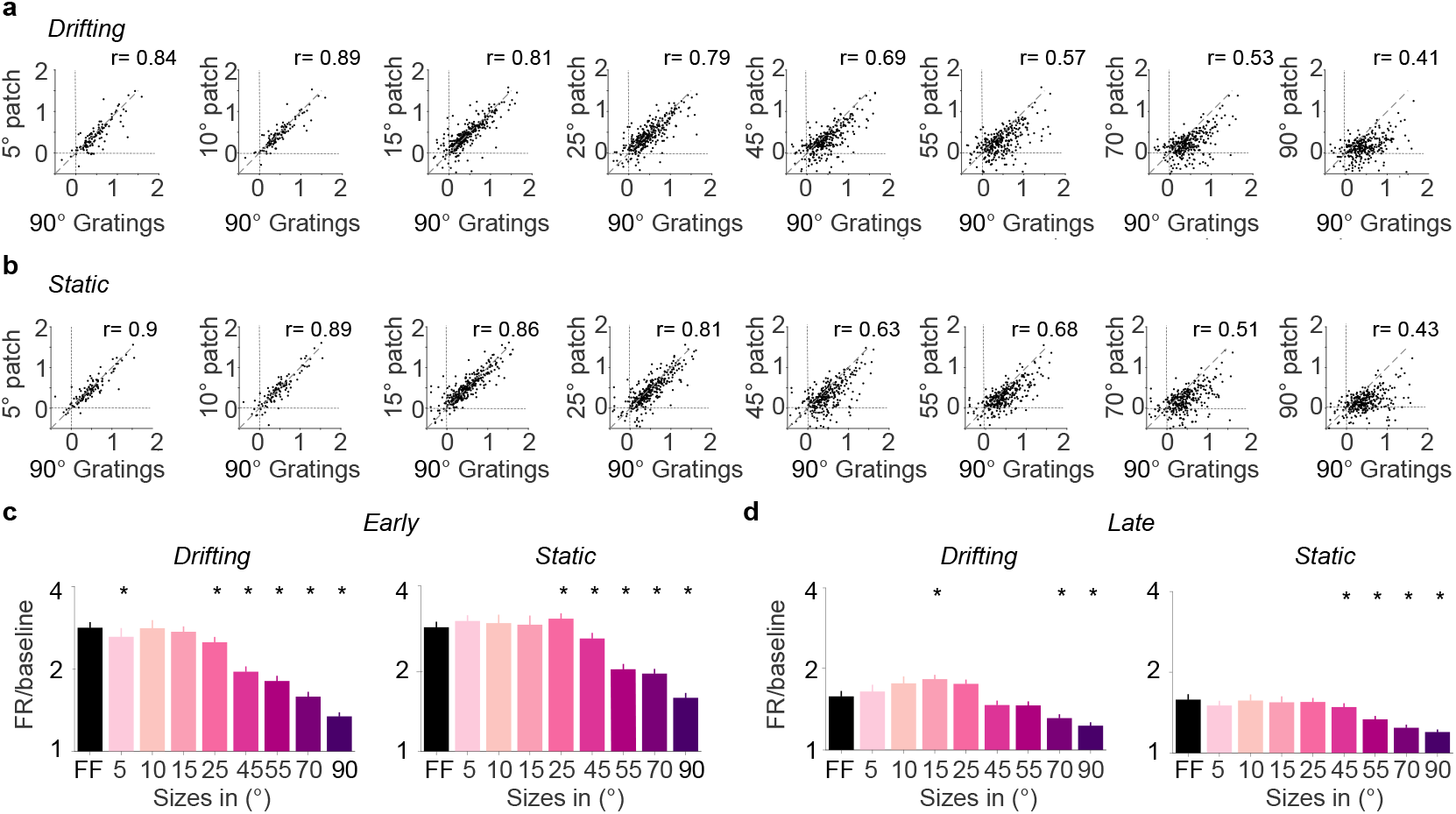
Firing rates to 90° gratings *vs* different sizes of the patches covering gratings. a) Scatter plots comparing 90° Gratings (we define it as full-field) to different sizes of the gray patch for the drifting condition (r-Pearson correlation coefficient). b) Same as in a for stationary conditions. c) Full-field gratings (90°) compared to different sizes of the gray patch covering gratings during the early stimulus presentation from 0.04 s to 0.15 s (Mean and the SEM. * p-values *<* 0.01 units per size 5° and 10°, *n* = 117 neurons, for sizes *>* 10°, *n* = 335 neurons, 6 animals). d) Same as c) but for the late stimulus period 0.2 s to 1 s.

**Figure S3.**
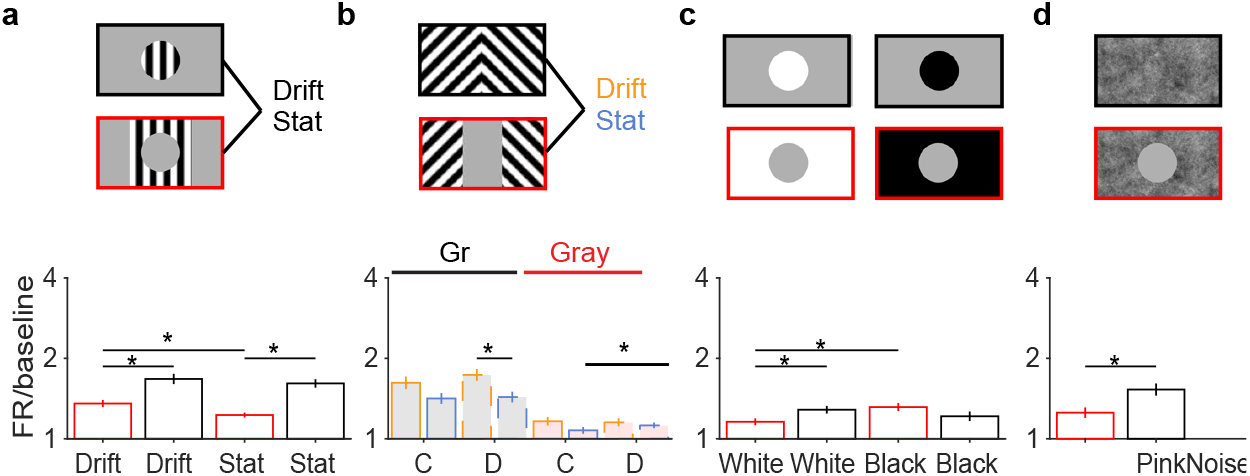
Comparison of firing rate during the late period for larger sizes for all protocols. a-d) Stimuli on the top, plots represent the average of the population firing rate during the late period from 0.2 s to 1 s, each unit is normalized to baseline, for sizes ≥ 45°. Each protocol includes different units and is normalized to the baseline for each block (*p-values *<* 0.01, Wilcoxon signed-rank test, each group is compared within the same protocol).

**Figure S4.**
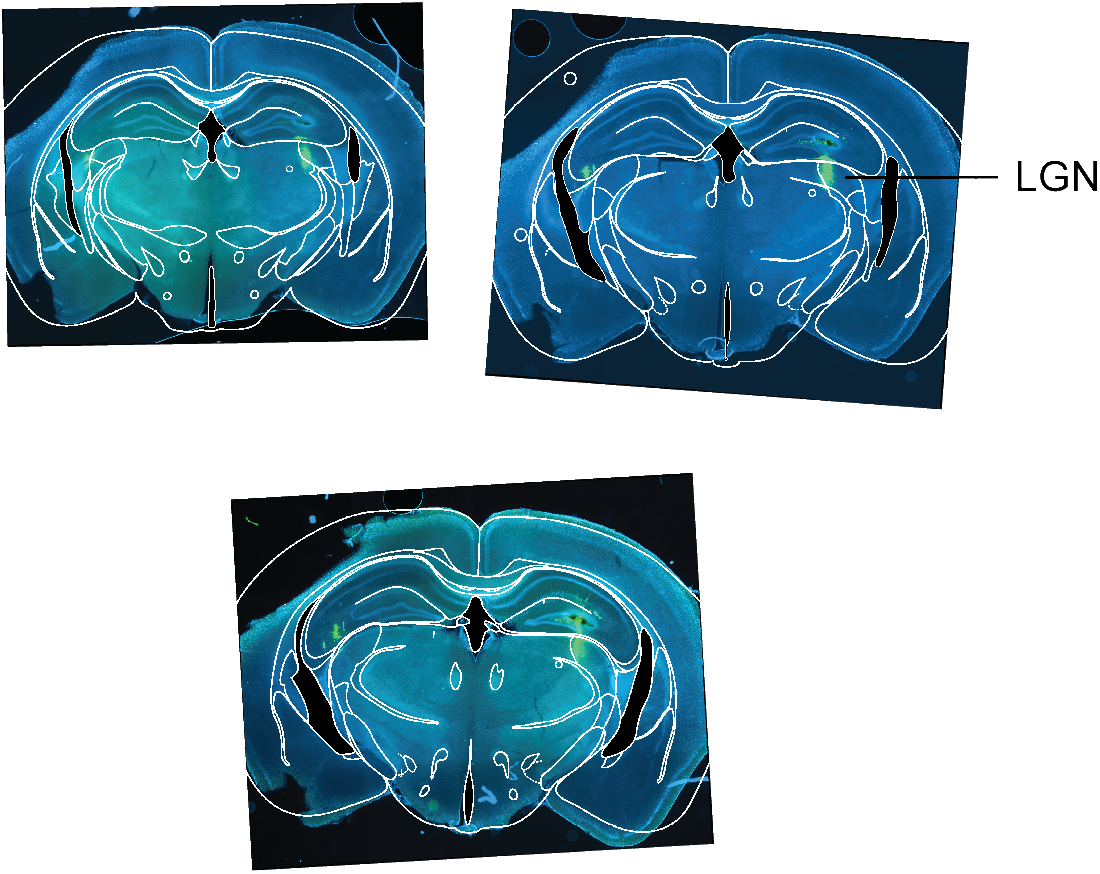
Histological confirmation of LGN recordings. Coronal sections of the mouse brain. During each recording session the Neuropixel probe was covered with DiD dye, this allowed to track of the recordings site at the end of each experiment. Histological sections of 100 *μ*m were observed under a fluorescent microscope. Representative images of one brain confirm that our coordinates targeted LGN (−1.94 mm, -2.06 mm and -2.18 mm relative to Bregma).

**Figure S5.**
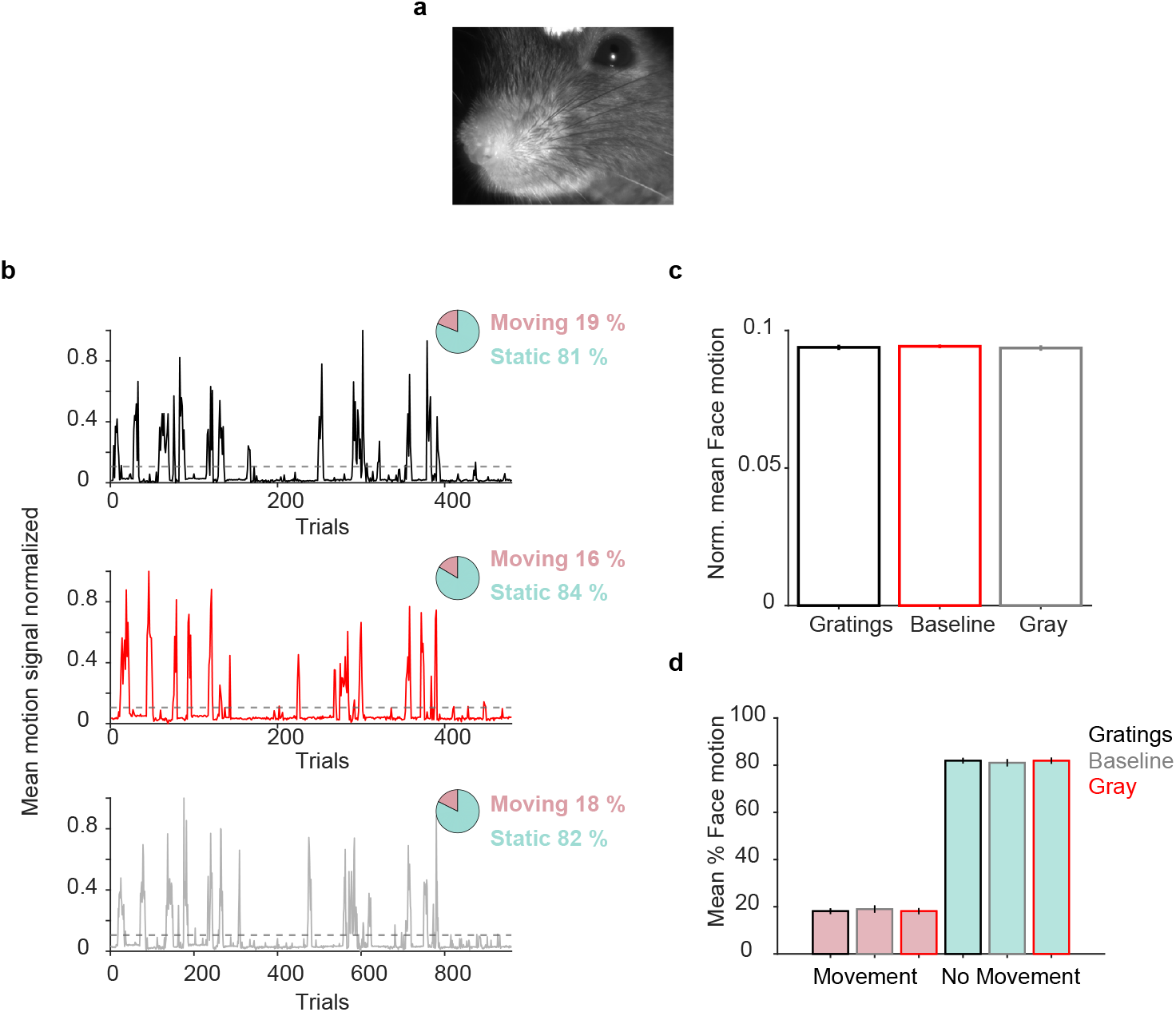
No difference in face movement during gratings, gray and baseline. a)Example of a ROI of a frame to analyze face movement. b) Example of the normalized motion signal during one session. The trials are separated into gratings, gray, and baseline, and the face movement is compared between conditions. The top right inserts represent the percentage of trials that crossed a threshold value (in dashed gray line). c) Comparison of the mean normalized face motion across sessions (20 sessions). Mean and SEM per session were then compared across sessions. d) Comparison of the percentage of movement and no movement across sessions. The percentages were calculated per session and averaged across sessions (20 sessions).

**Figure S6.**
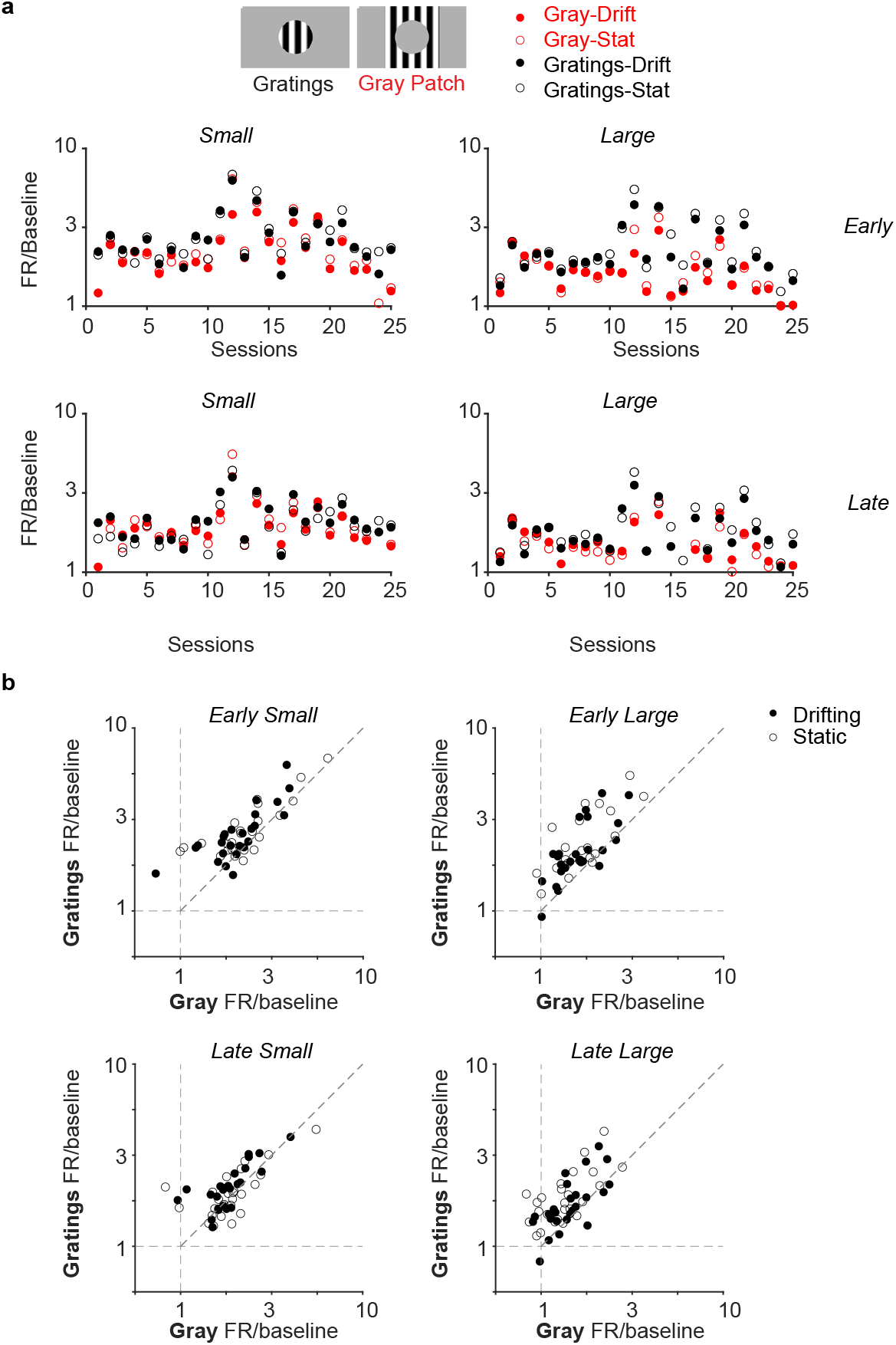
Analysis per session shows similar effects. a) Mean of each condition per session, divided into early (0.04 s - 0.15 s) and late (0.25 s - 1 s) stimulus periods. Each session has a different number of units. b) Scatter plots comparing the mean response per session between gratings condition and Gray patch condition, overall per session the responses are maintained as we previously presented comparing all units.

